# Effect of very long-chain lipids on the organization of biological membranes: A simulation perspective

**DOI:** 10.1101/2025.04.08.647797

**Authors:** A. Quas, C. Rickhoff, A. Heuer, R. Wedlich-Söldner

**Author notes:** Institute of Physical Chemistry, University of Münster (A. Quas).

## Abstract

Lipids in all biological membranes are distributed heterogeneously across the bilayer. A particularly striking example for this asymmetry is the yeast plasma membrane (PM), which exhibits a high concentration of very long-chain sphingolipids (SL) in its outer leaflet. Experimental observations indicate the existence of highly ordered gel-like PM domains that are enriched in SL but depleted in the major yeast sterol ergosterol. For a better mechanistic understanding of these unusual domains we have performed coarse-grained molecular dynamics simulations with membranes containing varying concentrations of very long-chain lipids. In agreement with experimental results we observed formation of a gel phase, with high order parameter of the acyl chains and hexagonal arrangement of lipid tails, at higher concentrations of very long-chain lipids. Our simulations also show that ergosterol is excluded from these gel regions and that, even when embedded into a liquid disordered phase, gels remain stable on the simulation time scale.

## I. INTRODUCTION

The different lipid species of a cell’s plasma membrane (PM) are distributed heterogeneously across the leaflets [1][2]. This asymmetry is essential for the organization and function of the PM and thus for the survival of the cell. In yeast *Saccharomyces cerevisiae*, sphingolipids (SL) account for about 10-20% of membrane lipids. They possess very long fatty acids (VLCFA), mainly fully saturated C26, and are concentrated in the outer layer of the plasma membrane.[3] Combining anisotropy and fluorescence lifetime measurements of packing-sensitive membrane probes it was proposed that SL organize into highly ordered gel domains in the yeast PM. Their structure and composition appears unique, as they mainly consist of SL and are depleted for sterols.[4] In contrast, liquid-ordered domains of animal cells are enriched in both, SL and cholesterol [5]. SL have also been associated with the transport, assembly and function of the major yeast H^+^-ATPase Pma1 [6]. This protein uses energy to transport proton out of the cell and, together with the vacuole ATPase maintains cytosolic pH. Via the protein gradient across the PM, Pma1 is also largely responsible for nutrient and small molecule uptake by yeast cells, which is facilitated by a large number of proton supporters.[7] Interestingly, recent cryo-EM measurements revealed that Pma1 forms hexametric oligomers in native membranes that surround a crystalline array of 57 lipids in the outer leaflet [8]. The apparent shape of these lipids in the EM images is consistent with saturated SL [9]. However, the precise identity of these lipids and a potential link to gel-like PM domains has not been examined. To adress these questions, it is important to study the fundamental influence of VLCFA on the properties of membranes. Investigating these questions experimentally has inherent limits. Fluorescence microscopy, for example, cannot achieve simultaneous high temporal and spatial resolution. Collective observation of multiple membrane components is also experimentally challenging.[10] A powerful alternative to understand the molecular mechanisms and underlying dynamics of membrane systems is molecular dynamic simulations. Despite significant computational advances over recent years, the long time scales and large system sizes required to simulate complex membrane mixtures remain a challenge for atomistic simulations. This has motivated the application of increasingly accurate coarse-grained models.[11] In this study, we performed coarse-grained simulations of yeast PM bilayers using the Martini2 force field [12]. We implemented lipid mixtures that closely resemble the native environment of the yeast PM. By varying the asymmetry and relative amounts of VLCFA in our simulations we then investigated the conditions that allow formation of gel domains or gel phases. To study inter-leaflet interactions of VLCFA, we compared simulations with asymmetric membranes (long-chain lipids exclusively in the outer leaflet) as well as symmetric membranes with longchain lipids in both leaflets. In addition, we characterized the lateral distribution and flip-flop dynamics of the yeast sterol ergosterol in either system.

## II. METHODS

### A. Systems

The chosen membrane composition was based on published values [13] and whole-cell lipodomics results obtained in the Wedlich-Söldner group. As there are currently no force field parameters for yeast sphingolipids we instead modeled SL by VLCFA-phosphatidylinositol (loPI) with one C18:0 and one C26:0 VLCFA (figure 1). Experimentally, this approximation is validated by findings that yeast cells require long-chain lipids but can substitute SL with loPI [3]. In addition, the head groups of PI and the common yeast SL, inositol-phosphoryl ceramide (IPC), have a similar chemical structure. Two different systems were considered: asymmetric membranes with loPI exclusively in the outer leaflet (l-s), symmetric membranes with loPI in both leaflets (l-l) or without any loPI (s-s). For the inner leaflet of the l-s systems, phospholipids with a chain length of 16/18 were selected to simplify the system (table I). The composition of both leaflets in the symmetric systems correspond to the composition of the asymmetric outer leaflet (l-l) or the inner leaflet (s-s). For all systems, each leaflet contained about 676 lipids, leading to a system size of about 18 nm x 18 nm. All membrane systems were set up with identical numbers of lipids in either leaflet. Although diverging area per lipid values are expected for the inner and outer leaflet of the l-s systems, we assume equilibration of the systems by flip-flop of ergosterol within the first microsecond (see figure 17). Simulations were performed at temperatures ranging from 283 K to 313 K with an interval of 10 K.

**Figure 1.**
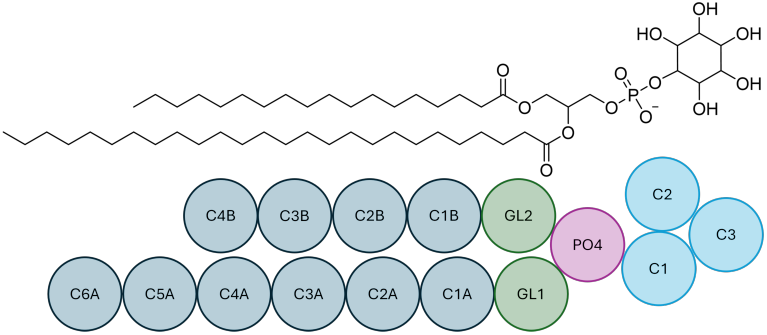
Structure of the long-chain lipid loPI modeling the long-chain sphingolipids of the plasma membrane.

**Table I.**
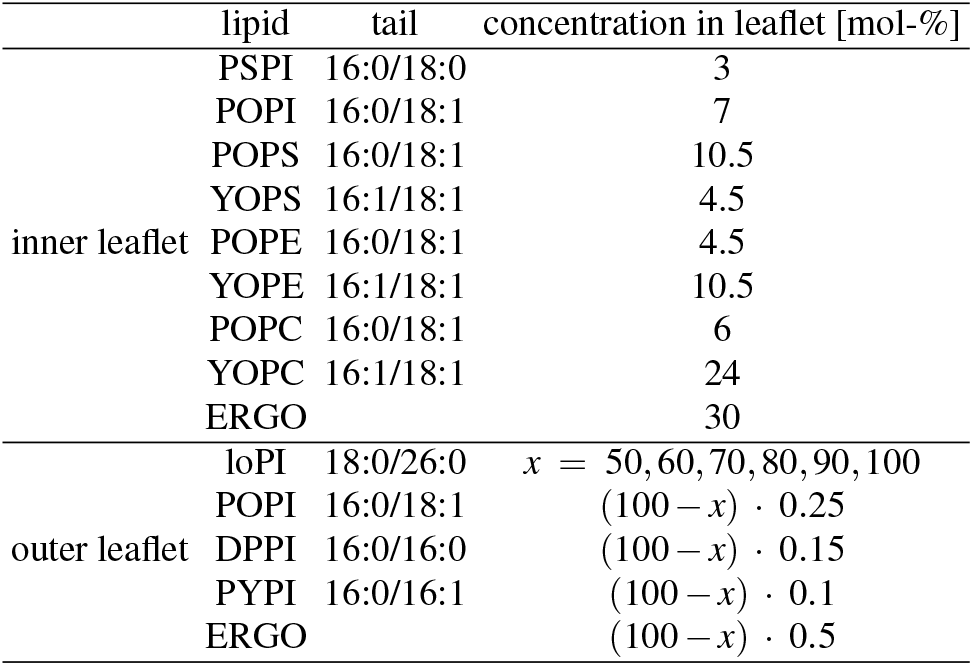
The composition of the inner and outer leaflet of the simulated l-s membranes. The long-chain PI that models the long-chain lipids is abbreviated as loPI.

To specifically study the dynamic behavior of ergosterol, a three-component symmetrical bilayer was generated with loPI (260 molecules per leaflet) on the left, ergosterol in the center (130 molecules per leaflet) and POPI (260 molecules per leaflet) on the right. This simulation was performed at 303 K for 4 *µ*s.

### B. Simulation details

The membrane-systems were set up with the membrane builder *insane* [14] and simulated with Gromacs 2019.6 [15][16] using the Martini2 force field [12]. Martini2 instead of Martini3 was chosen because the latter mainly focuses on improvements in protein interactions, which are not of relevance for this project [17][18]. The known differences to atomistic simulations due to coarse-graining, such as different melting points, must be taken into account in both models. We therefore expect similar results for both Martini versions.

The equilibration process consisted of five steps, in which the time step was increased from 2 fs to 20 fs and the position restraints on the lipids reduced. The velocity-rescale thermostat [19] and the Berendsen barostat [20] were applied. For the production simulations, the time step was set to 20 fs. The temperature was controlled by the velocity-rescale algorithm [19] and the Parrinello-Rahman [21] barostat with semi-isotropic pressure coupling was applied to control the pressure (1 bar). The lipid-bilayer and the solvent molecules were treated in separate temperature and pressure coupling groups. The coulomb interactions were calculated based on reaction field electrostatics with a cut-off of 1.1 nm, while a plain cutoff of 1.1 nm with potential-shift-verlet modifier was used for the evaluation of the Van der Waals interactions. The systems were simulated for 2 *µ*s (symmetric membranes) or 4 *µ*s (l-s membranes). The first microsecond of the production run was classified as equilibration to ensure that the ergosterol concentrations in both leaflets of the l-s system had also reached equilibrium.

Most analysis was performed with the python library MD-Analysis [22]. VMD [23] was employed to create the snapshots of the systems.

### C. Analysis methods

Different parameters were evaluated to characterize the membrane systems. To describe the lipid neighborhood, the hexagonal order parameter 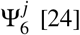, with *θ* being the angle between the *x*-axis and the vector connecting lipid *j* with its neighbor *m*, was calculated. If the lipid *j* is perfectly hexagonally surrounded, 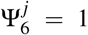. *τ*_*α*_ was chosen to be 1 ns, as the variance was relatively constant with respect to the parameter at values higher or equal 1 ns (see figure 18)

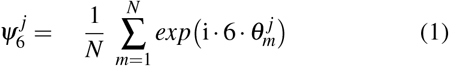

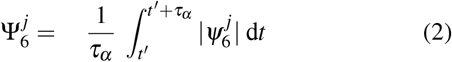

The order parameter describes the alignment of the lipid in relation to the membrane normal. The angle *θ* is defined as the angle between the membrane normal and the vector connecting two consecutive coarse-grained beads. The brackets indicate ensemble and time average. The parameter ranges from 1 (straight tail along the membrane normal) to -0.5 [25].

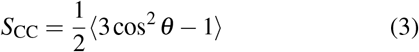

The coupling of the leaflets significantly depends on the number of atoms available for interactions in the area of overlap. Introducing a mean-field character, the bead-density is used as a measure for the number of interaction sites. Assuming that the lipid density of the upper leaflet at position *z* is significantly lower than that of the lower leaflet, increasing the density of the lower leaflet would not increase the number of inter-leaflet interaction sites. Therefore, the number of interaction sites is limited by the minimal density *ρ*_min_(*z*).

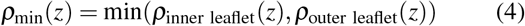

Integration along the membrane normal *z* yields a measure of interdigitation that is proportional to the number of actual interactions of lipid fragments of lipids belonging to opposite leaflets:

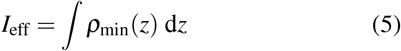

## III. RESULTS

### A. Phase behavior

As a first step to investigate the membranes’ properties, we characterize the phase behavior. By visual inspection of the snapshots of the l-s systems (see figure 2 top), the two leaflets show major differences. The lipids in the inner leaflet exhibit a disordered structure in both loPI-concentrations shown. On the other hand, the lipids in the outer leaflets have a well-defined structure and straight lipid tails at high concentration of loPI. Comparing the l-s system with 90 mol-% loPI in the outer leaflet with the l-l system, the same observations can be made for both leaflet at 283 K (see figure 2 bottom). At 313 K, the lipid structure is no longer visible and the membrane thickness is reduced compared to the system at 283 K. The membrane is significantly more compressed.

**Figure 2.**
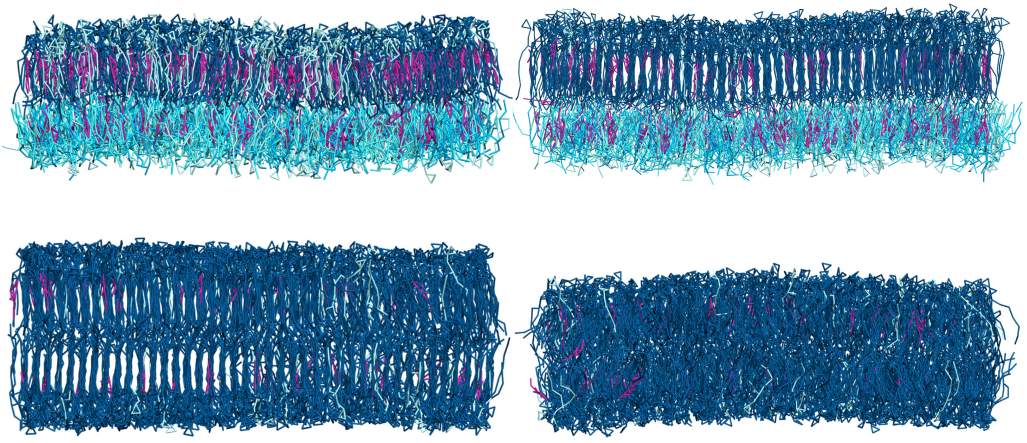
Snapshots of the l-s membranes at 283 K with 50 mol-% loPI (top left) and 90 mol-% loPI (top right) and of the l-l membranes with 90 mol-% loPI at 303 K (bottom left) and 313 K (bottom right). The long-chain lipids loPI are shown in dark blue, ergosterol in pink and other phospholipids in light blue.

We calculated the hexagonal order parameter 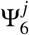 (equation 2) for the systems to characterize the structure of the neighborhood. Lipids in the gel phase typically take on a hexagonal structure, resulting in values close to 1 for this parameter. Plotting the lipid tail position and the corresponding value of 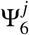 , differently structured areas in the membrane can be identified (figure 3). The outer leaflet of the l-l system with 50 mol-% loPI at 283 K shows well structured regions with high values for 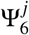, but also regions without a hexagonal arrangement resulting in lower values. Increasing the loPI concentration to 100 mol-%, most of the lipids in the leaflet are in a hexagonal neighborhood structure. For both concentrations shown, the hexagonal structure of the lipids is not observed at 313 K. This increase in the hexagonal ordering in the l-l systems at temperatures lower than 313 K is also observed for all investigated temperatures in the l-s membranes (figure 4 top). Here, the parameter increases from about 0.62 at 50 mol-% loPI to about 0.76 at 100 mol-% loPI at 283 K indicating an increasing crystallinity in the membrane. Lower temperatures result in a higher hexagonal order parameter for all concentrations due to the lower entropy. The order parameter *S*_CC_ describing the lipid tail alignment along the membrane normal (equation 3) shows the same profile as 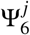 (figure 4 bottom). The last two beads (C5A and C6A) were excluded from the calculation due to their deviating behavior (figure 7). The temperature dependency of the order parameter is more pronounced at lower concentrations. At 50 mol-%, the difference between 283 K and 313 K is about 0.13, while it is about 0.03 in the l-s systems with 100 mol-% loPI. The stronger binding between the lipids at higher concentrations results in a weaker temperature dependence of these systems. The high values of about 0.9 in the l-s systems with 100 mol-% loPI reflect, that the lipid tails are oriented parallel to the membrane normal. Together with the hexagonal arrangement of the lipid tails, the alignment of the lipid tails indicate that the membranes are in a gel phase at high concentration of loPI. The obtained values are in agreement with experimentally determined acyl chain order parameters of lipids in the gel phase [26].

**Figure 3.**
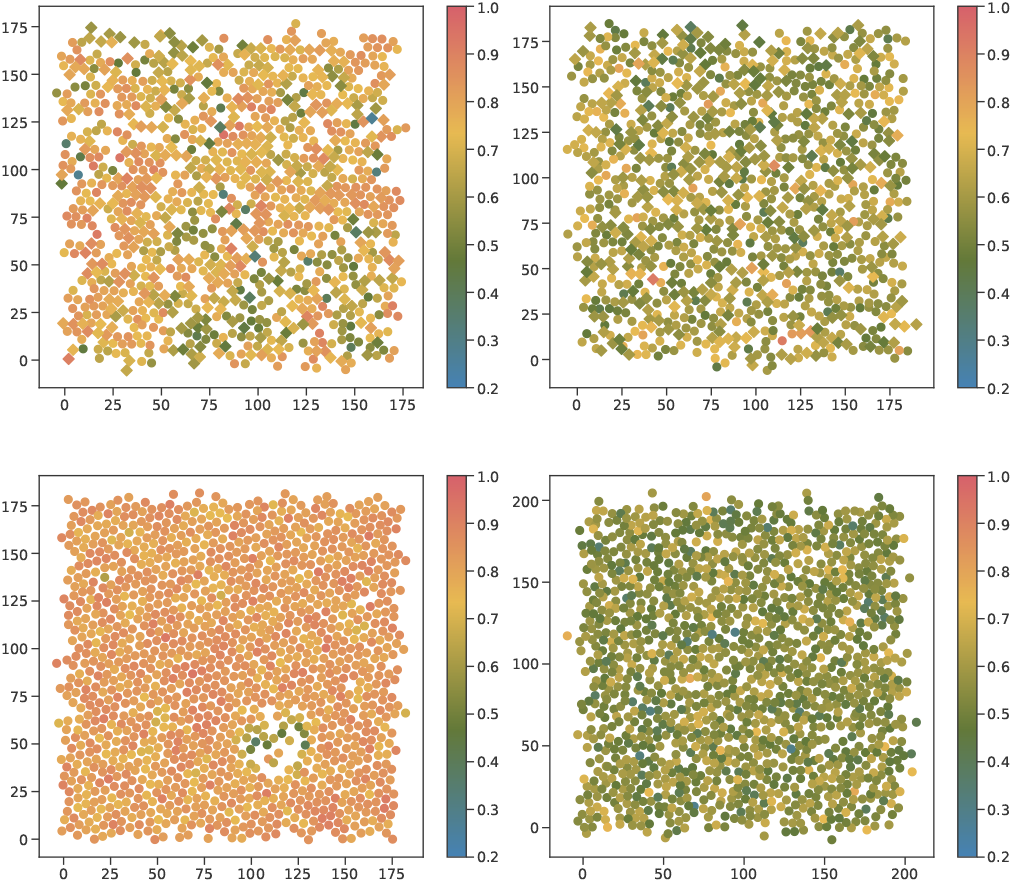
View on the *xy*-plane of the membrane, where each point corresponds to a lipid tail. loPI lipids are shown as circles, other phospholipids as squares. The color represents the value of the hexagonal order parameter 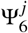. The beads C2A and C2B were chosen for calculation. The upper leaflet of the l-l membranes with 50 mol-% (top) and 100 mol-% loPI (bottom) at 283 K (left) and 313 K (right) are shown.

**Figure 4.**
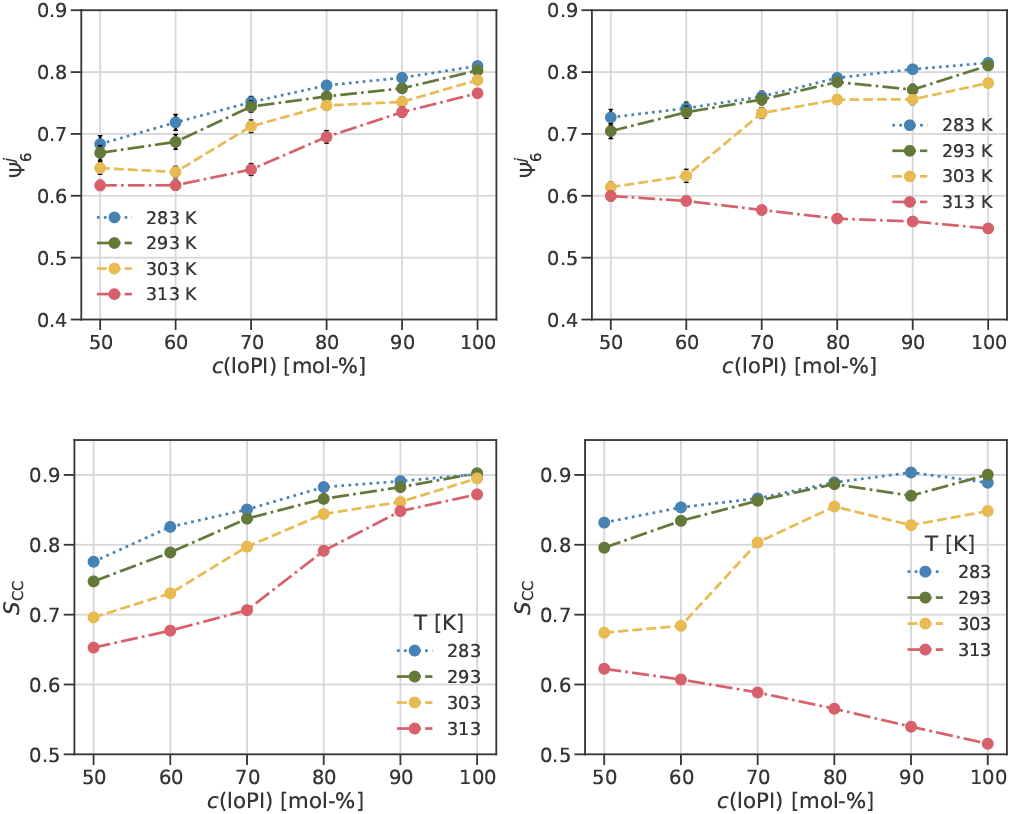
The mean hexagonal order parameter 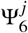 and the order parameter *S*CC for the l-s membranes (left) and l-l membranes (right) for varying concentrations of loPI. The error bars of 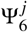 correspond the mean variance of the parameter within the membrane.

In contrast to the l-s systems, the order parameter of the l-l membrane decreases at 313 K from about 0.62 at 50 mol-% to 0.51 at 100 mol-% loPI. This reflects a transition from the gel to the liquid phase in the l-l membranes below this temperature. As the concentration of loPI increases, the ergosterol content decreases (*c*(ERGO) = (100 −*c*(loPI)) ·0.5), explaining the decrease in the order parameter for the system at 313 K due to the ordering effect of ergosterol [27].

Experimentally, the melting point corresponding to the phase transition from gel to liquid of IPC with the same chain length as loPI was determined by DPH fluorescence anisotropy [28]. To compare the resulting value of 327 K, the temperature shift due to the coarse-graining has to be taken into account. For DPPC, a decrease of 40 K for coarse-grained as compared to atomistic was found [29]. Assuming the same shift for the l-l membranes investigated in this project, the melting temperature is expected to be 303 K + 40 K = 343 K and 313 K + 40 K = 353 K. Whether this assumption is correct and whether the head group has a major influence on the melting temperature needs to be investigated further. A priori it is not clear whether only the long-chain lipids build up the hexagonal structure and the short-chain lipids appear to be defects within the crystal structure of the gel, or whether the short-chain lipids also adopt a hexagonal arrangement due to the structuring of the long-chain lipids. Calculating 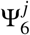 for long-chain and short-chain lipids separately reveals, that the parameter also increases for the short-chain lipids and is about 0.03 smaller than for the long-chain lipids for all investigated concentrations of loPI (figure 5 left). This indicates, that the short-chain lipids adapt the structuring of the long-chain lipid and that they are included into the gel phase. The mixing of long and short-chain lipids can also be quantified by the fraction of loPI neighbors around loPI (figure 5 right). Assuming a perfectly mixed leaflet, the fraction of neighbors is identical to the concentration of loPI. To account for ergosterol molecules, that might have flipped into the outer leaflet, the loPI concentration in equilibrium is considered. For all concentrations and temperatures, the fraction of loPI-loPI neighbors is linearly increasing with the loPI concentration. The values obtained are slightly higher but close to the reference of a random neighbor distribution (figure 5 right, grey line). loPI-loPI neighbors are therefore slightly preferred, but there is no separation of the long-chain from the short-chain lipids as already indicated by similar high values for the hexagonal order parameter.

**Figure 5.**
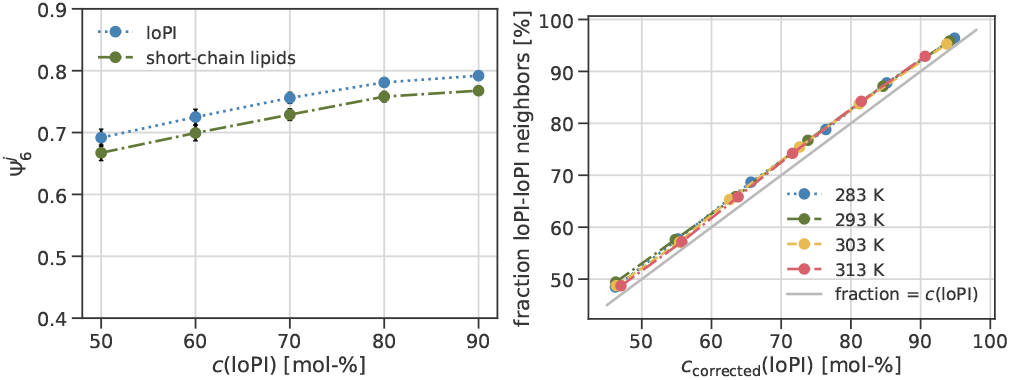
The mean hexagonal order parameter 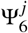 for the l-s membranes at 283 K for loPI and the short-chain lipids in the outer leaflet (left). The fraction of loPI neighbors around loPI within a radius of 12.17 Å at different concentrations of loPI corrected for the flipping of ergosterol for the l-s membranes.

The order parameter *S*_CC_ is also calculated for the phospholipids of the inner leaflet of the l-s systems and compared to the s-s membranes to investigate whether the long-chain lipids in the outer leaflet influence the ordering in the opposing leaflet (figure 6). Negative values of Δ*S*_CC_ represent a higher lipid order in the s-s system compared to the inner leaflet of the l-s system. The results show, that the lipids in the inner leaflet of the l-s system are less ordered than in the s-s system with the same composition. The difference is more pronounced at higher temperatures. The highest difference is observed at 313 K for the system with 80 mol-% loPI in the outer leaflet. Here, the order parameter is about 0.08 higher in the corresponding s-s system. This shows an influence of the long-chain lipids in the outer leaflet on the ordering of the lipids in the opposing leaflet. We suppose, that the reduction in ordering as compared to the s-s system aims to stabilize the gel phase in the outer leaflet. Lipids with a lower order parameter are more flexible in their conformation and can therefore adapt more freely to the rigid gel phase on the other side.

**Figure 6.**
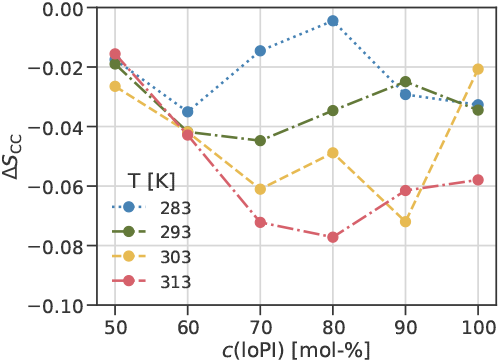
The difference of the order parameter of the phospholipids in the inner leaflet of the l-s system and the s-s system with the same lipid composition at the same temperature.

### B. Lipid structure

Lipids with a high asymmetry in the tail lengths are found in different conformations in membranes: interdigitation into the opposing leaflet is observed as well as bending towards the headgroup.[30] We therefore analyzed the orientation of each tail section of the longer tail of loPI with the order parameter (figure 7). The tail part close to the headgroup exhibits the highest order parameters for both the l-s and the l-l membranes. Values higher than 0.8 reflect a straight lipid tail in this region. Going down the tail, the order parameter decreases. For the beads C4A and C5A, the order parameter reduces to about 0.5 at 313 K. The lowest ordering is found for the last part of the tail. Due to the phase transition, the order parameters of all tail parts of the l-l systems decline at 313 K. The reduction in order is less for the last part of the tail, as this part is also not well aligned in the gel phase. A smaller temperature dependence is also observed for C5A-C6A in the l-s system compared to the other tail parts. Intuitively, one would assume straight tail ends to facilitate interdigitation into the other leaflet for longer tails. However, an order parameter close to zero suggests an almost random alignment with the *z*-axis. It is surprising that the long tail is needed for gel formation, but the long tail end itself is not part of the gel phase. The behavior of the last part of the tail is further analyzed (figure 8). For the l-s system, the order parameter increases linearly from 0.1 at 50 mol-% loPI to 0.24 at 100 mol-% loPI. No significant influence of the temperature is observed for these systems. In the l-l systems, the order parameter increases up to 0.35 in the investigated concentration range for temperatures lower than 313 K. The order parameter is therefore about 0.16 higher in the l-l membrane at 100 mol-% loPI compared to the corresponding l-s membranes. Although these values are significantly smaller than for the beads close to the headgroup, the higher ordering in the l-l systems could be due to a stronger interdigitation caused by the presence of long-chain lipids in both leaflets. At 313 K the increase in the order parameter is smaller for the l-l system compared to lower temperatures. This trend is in contrast to the rest of the tail whose order parameter decreases with increasing concentration (figure 4 right). Comparing the order parameter of the last bead to the behavior of the membrane thickness, similarities become apparent. Increasing the loPI concentration leads to a nearly linear thickening of the l-s membranes. If the headgroups are further apart, the long tail has the space to align itself, which results in a higher order parameter. Since both leaflets in the l-l system contain long-chain lipids, which increase the membrane thickness with increasing concentration, the thickness and its increase is larger at temperatures lower than 313 K. Because the lipid tails are not aligned with the *z*-axis but unstructured in the liquid phase, the membrane thickness increases considerably less at 313 K in the l-l membrane. This creates less space for the long tail end to straighten up, resulting in a smaller increase in the order parameter.

**Figure 7.**
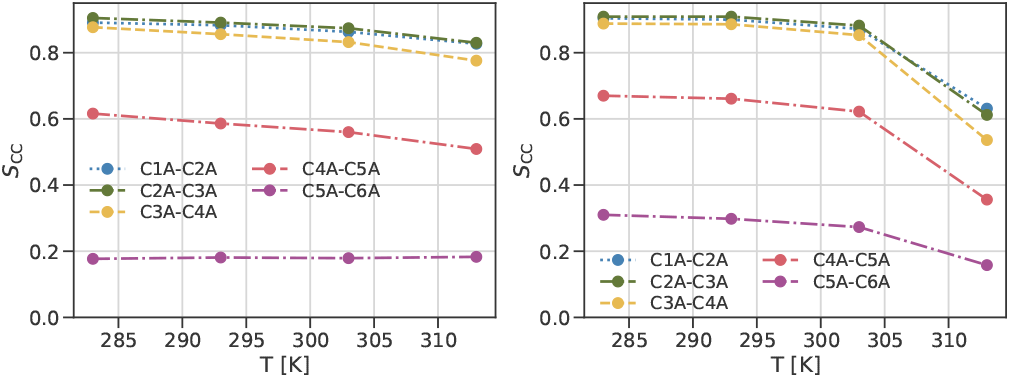
The order parameter for neighboring beads in the long tail of loPI at different temperatures and the system with 80 mol-% loPI. The results for the l-s membranes are shown on the left side, for the l-l systems on the right.

**Figure 8.**
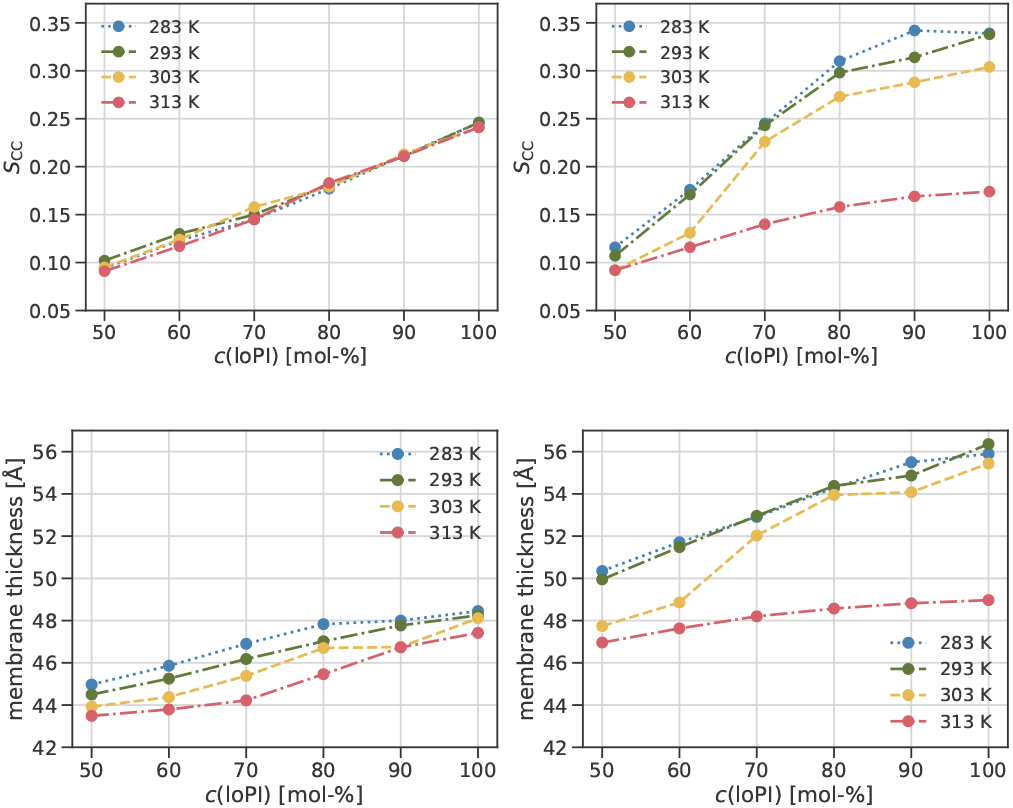
The order parameter *S*_CC_ (top) of the last two beads (C5A and C6A) in the long tail of loPI and the membrane thickness (bottom; the PO4 beads are used as reference) in the l-s (left) and l-l systems (right) at different concentrations of loPI.

**Figure 9.**
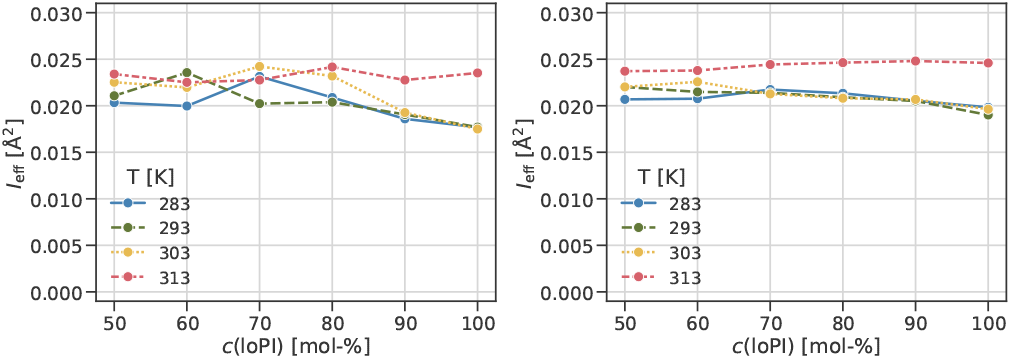
The interdigitation *I*_eff_ depending on the concentration of loPI at different temperatures for all phospholipids and ergosterol in the l-s (left) and l-l membranes (right).

### C. Inter-leaflet coupling

The analysis of the phase behavior revealed a lower melting temperature of the l-l membrane compared to the l-s system. This is probably due to deviating inter-leaflet interactions. Assuming that the presence of beads of both leaflets at the leaflet interface region causes the coupling, integrating the minimal density of both lipids along the membrane normal results in a parameter proportional to the number of sites for inter-leaflet interactions. Taking into account all lipid-types including ergosterol, similar values for *I*_eff_ can be observed for the l-l and l-s membranes. It is remarkable that the interdigitation *I*_eff_ is significantly higher at 313 K and high concentrations of loPI not only for the l-l but also for the l-s system. For the l-l system, the phase transition to the liquid phase explains the deviating trend for this temperature as the lipid structure significantly changes leading to higher values of *I*_eff_. As the phase transition is not observed in the l-s system, the interdigitation does not seem to be the primary source for this behavior. Additional insight about the interdigitation can be found in the supporting information.

### D. Behavior of ergosterol

Ergosterol is a main component of the plasma membrane and has a significant influence on its properties. Here we investigate how ergosterol behaves in the gel phase and whether our simulations confirm a lower ergosterol content as found experimentally [4].

The flip-flop rates of ergosterol are calculated for the l-l membranes (figure 10). As expected, the rates in the systems increase with temperature. At temperatures lower than 313 K, a decrease in the flip-flop rate is observed. After the transition to the liquid phase, the flip-flop rate does not decrease with higher concentrations of loPI. This shows, that not the addition of long-chain lipids but the formation of the gel phase inhibits the flipping of ergosterol. A reason for this might be a blocking effect of the last tail part in the gel phase, as the low order parameter indicates bending of this part in the inter-leaflet area. These results can be transferred to the experimentally observed gel domain in the outer leaflet of the plasma membrane. Assuming that both the flow in and out of a leaflet in the gel phase is more difficult, this means that ergosterol molecules have difficulty flowing from the inner, liquid leaflet of the plasma membrane into the gel phase in the outer leaflet. Consequently, flipping processes in this direction should not influence the ergosterol concentration in the outer leaflets gel phase significantly. Not only flipping processes but also diffusion of ergosterol from the surrounding liquid phase can influence their ergosterol concentration within the gel domain. The behavior is analyzed by simulating a l-l system with a gel phase of loPI on the left side, a liquid phase of POPI on the right side and ergosterol molecules in between (see figure 11). From the snapshots of this system, the hexagonal structure of the gel phase can be seen. At the end of the simulation the gel phase is dissolved and all lipids are mixed in a liquid phase. The dissolving of the gel phase also shows in the neighbors of loPI. During the first 3 *µ*s, loPI is mainly surrounded by loPI, the number of POPI neighbors is significantly lower (figure 12). After this point, the values equalize indicating a homogeneous mixing of the two lipid types.

**Figure 10.**
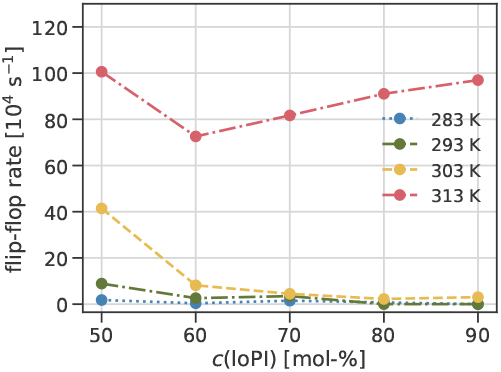
The flip-flop rate of ergosterol calculated for the l-l membranes for different concentrations of loPI.

**Figure 11.**
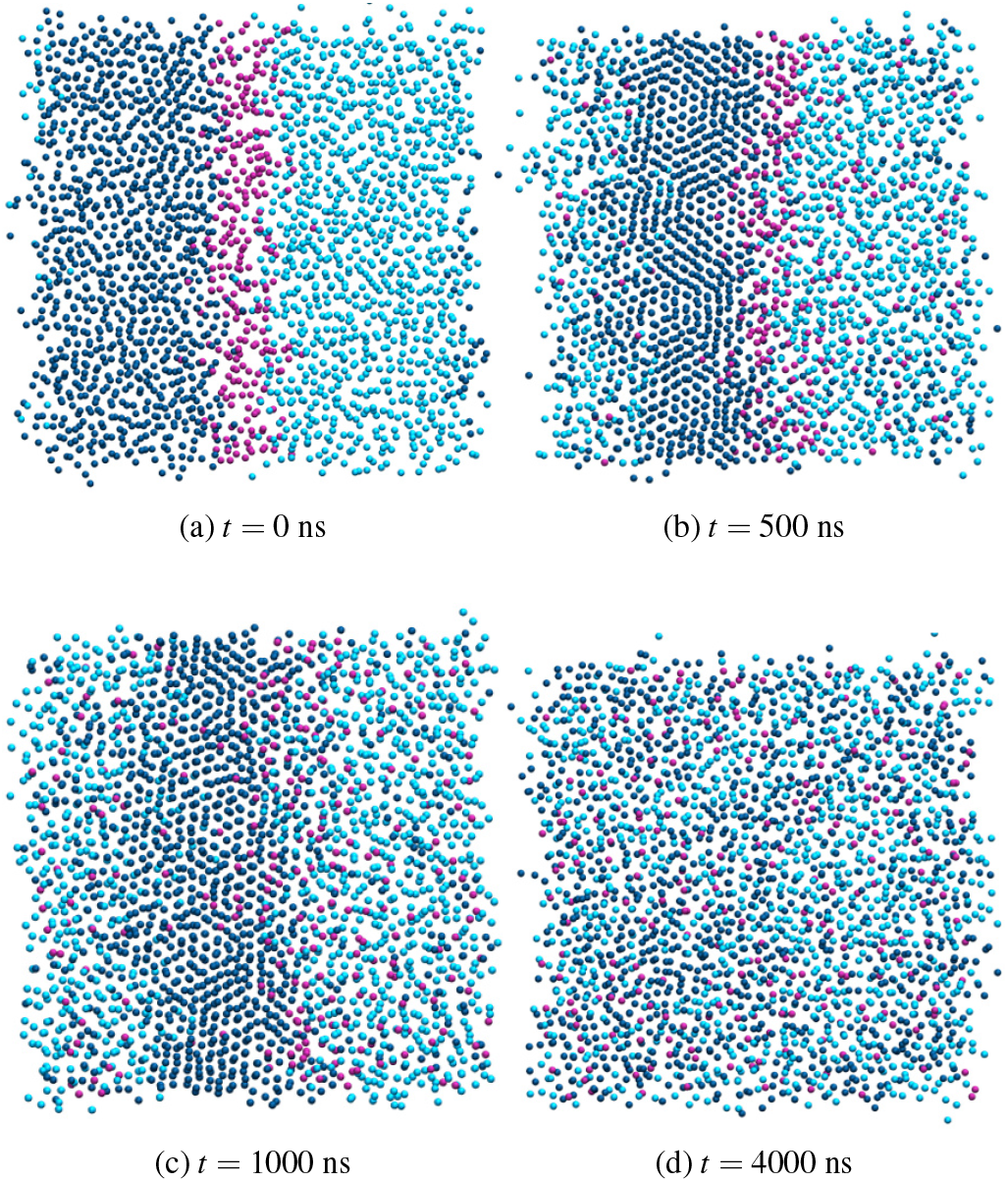
Top view of the membrane at different simulation times with the starting composition of loPI on the left side (dark blue), ergosterol in the middle (pink) and POPI on the right side (light blue). The beads C3A and C3B are shown for loPI and POPI and the ROH beads of ergosterol for better visibility.

**Figure 12.**
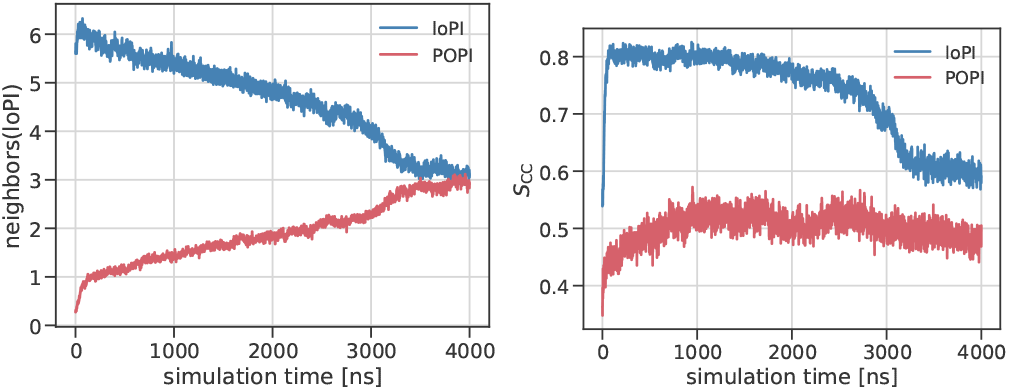
The average number of neighbors around loPI within a radius of 11.8 Å (left) and the order parameter of loPI (without beads C5A and C6A) and POPI (right) depending on the simulation time in the system with the starting composition of loPI on the left side, ergosterol in the middle and POPI on the right side (left).

The order parameter of loPI also sharply decreases at about 3 *µ*s displaying a phase transition from gel to liquid. The order parameter of POPI also slightly decreases at this simulation time. The average number of neighbors of ergosterol was calculated for the entire simulation time (figure 13). As expected, at the beginning of the simulation time ergosterol is primarily surrounded by other ergosterol molecules. As this ergosterol phase in the middle of the membrane dissolves during the simulation, the number of ergosterol in the neighborhood of ergosterol decreases. At the same time, the number of loPI and POPI around ergosterol increases as it diffuses into the two phases. It is noticeable that the number of POPI neighbors around ergosterol is higher than loPI, which is particularly pronounced at around 1000 ns. Starting from about 1.5 average neighbors, the number of POPI neighbors around ergosterol doubles during the first 500 ns, while the number of loPI neighbors only increases by a factor of about 1.3. This difference between POPI and loPI decreases as the simulation continues. After about 3000 ns, the number of POPI and loPI neighbors around ergosterol is about the same. This reflects the point at which the gel phase dissolves and the lipid types mix into a single liquid phase, as also shown by the order parameter of loPI (figure 13 right). The higher number of POPI neighbors indicates that ergosterol diffuses preferentially into the liquid phase. The free energy for going into the gel phase seems to be less favorable than going into the liquid phase. Together with the low flip-flop rate, the reduced diffusion into the gel phase makes a low ergosterol concentration in the gel phase very plausible.

**Figure 13.**
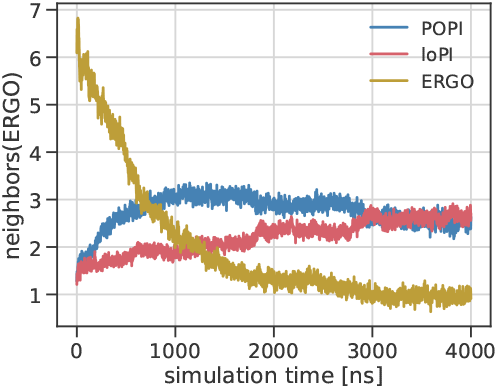
The number of neighbors of ergosterol within a radius of 11.5 Å depending on the simulation time in the system with the starting composition of loPI on the left side, ergosterol in the middle and POPI on the right side.

### E. Simulation of a gel domain

The simulations presented in previous chapters aimed to model the lipids’ behavior within a gel domain in the plasma membrane. To investigate the stability of a gel domain in a liquid environment, we simulated an asymmetric membrane with both a gel and a liquid phase in the outer leaflet. The lipid composition of this system is the same as for the l-s membrane with 60 mol-% loPI in the outer leaflet (see table I) but the loPI lipids are separated from the other phospholipids (see figure 14). The hexagonal order parameter was determined to evaluate whether the gel domain is intact at different simulation times (figure 15). At the beginning of the simulation, all loPI lipids are found on the left side of the membrane but only parts of them are hexagonal arranged as indicated by high values of the hexagonal order parameter. After 1000 ns, all loPI lipids belonging to the domain are in the gel phase. It can be seen, that some loPI diffuses into the other phase during the simulation time. The domain with high values for 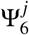 remains stable and is still observed after 4000 ns of simulation time. The lipid mixing is quantified by calculating the average numbers of neighbors around loPI (figure 16) (left). During the entire simulation time, the number of loPI neighbors is about six times higher than the number of short-chain neighbors. This shows that the loPI cluster forms a relatively stable gel domain. Due to diffusion of some loPI into the other phase, the number of loPI neighbors decreases and increases for short-chain lipids. This is also reflected in the order parameter of loPI. In the first nanoseconds, the gel domain is forming, resulting in an increase in *S*_CC_ to about 0.85. loPI diffusing out of the gel domain exhibit lower tail order and therefore lead to a decrease in the average order parameter. The evaluated parameters show that the gel domain is still intact after 4 *µ*s. The diffusion of some loPI lipids out of the gel domain suggests, that the composition of the membrane might not be the optimum. Starting with the same number of lipids in both leaflets might also influence the domain stability as it leads to flipping of ergosterol into the outer leaflet further increasing its concentration by more than 10% (see figure 26). The short-chain lipids surrounding the gel domain have an average order parameter between 0.6 and 0.7. Compared to the l-l system at 313 K (figure 4), the lipid tails are more aligned with the membrane normal. Together with an ergosterol concentration of more than abut 50 mol-% in this phase signifies that the gel domain is surrounded by a liquid-ordered phase.[31]

**Figure 14.**
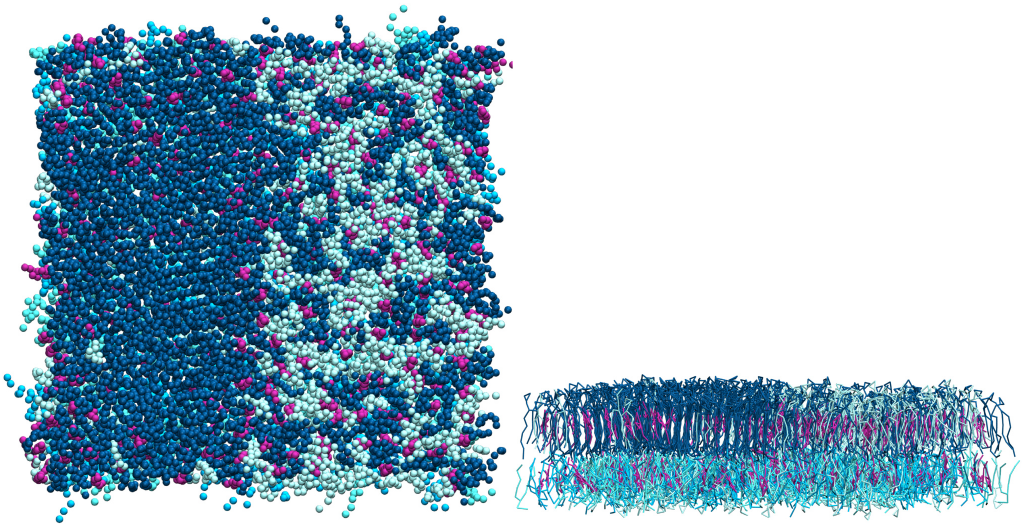
Snapshot of the simulated gel domain from top view (left) and side view (right) after 4 *µ*s. Ergosterol is shown in pink, loPI in dark blue and other phospholipids in light blue.

**Figure 15.**
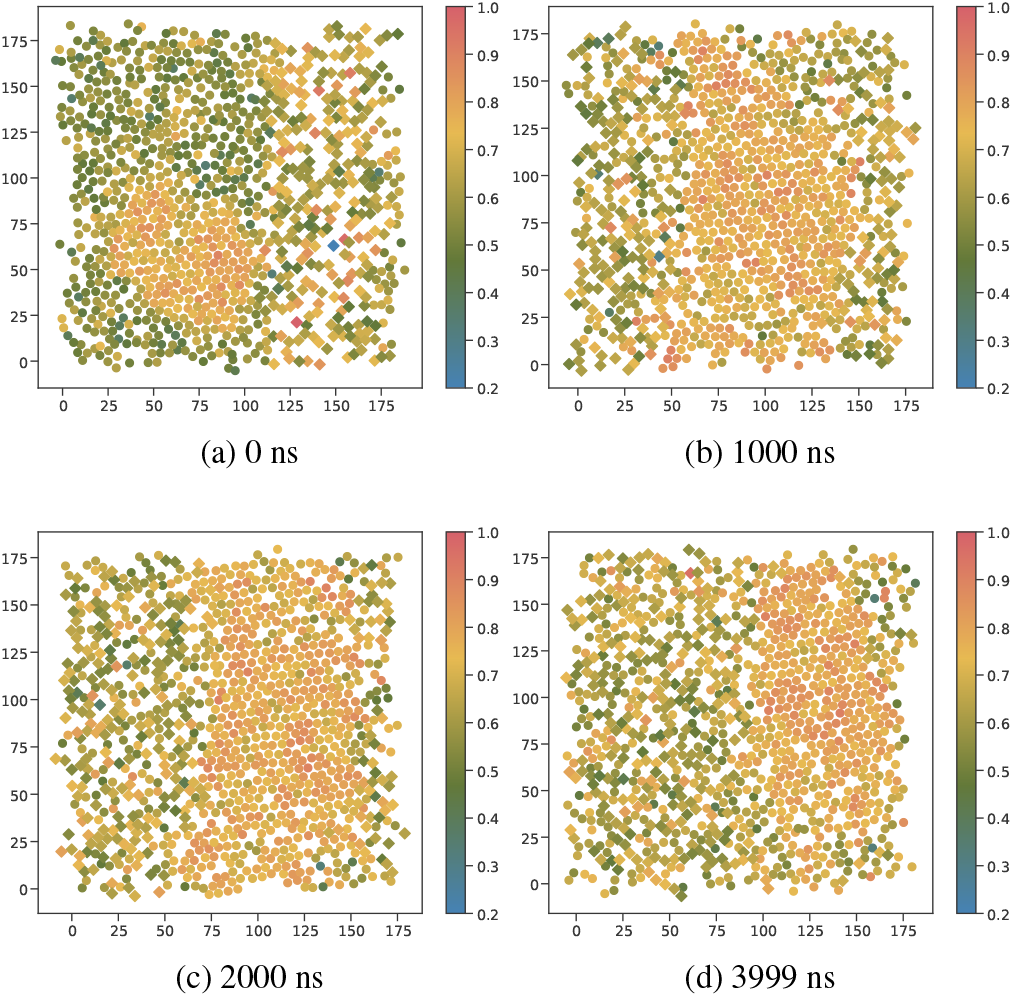
The hexagonal order parameter of the outer leaflet of the simulated gel domain at different simulation times. loPI is shown as circles, other phospholipids as squares.

**Figure 16.**
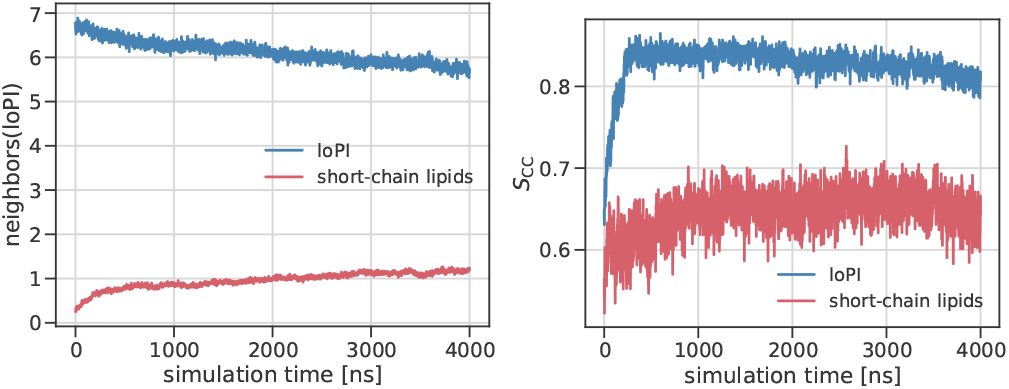
The average number of neighbors within a radius of 12.17 Å around loPI divided into loPI and short-chain lipid neighbors (left) and the order parameter *S*_CC_ of loPI without beads C5A and C6A and of the short-chain lipids as a function of the simulation time.

## IV. CONCLUSION

Experimentally, highly ordered gel domains with a high concentration of long-chain sphingolipids are found in the plasma membrane of yeast. We performed coarse-grained simulations of membranes with varying concentration of longchain lipids to investigate their impact on the membrane properties. A model lipid with the PI head group was used to simulate the sphingolipids.

Our analysis showed the formation of a gel phase at higher concentrations of long-chain lipids that is defined by an alignment of the lipid tails along the membrane normal and a hexagonal neighborhood structure. The end part of the long chain is not part of the gel phase structure and shows less alignment along the *z*-axis. In contrast to the l-s membranes that contain long-chain lipids only in the outer leaflet, the l-l system with these lipids in both leaflets show a transition from the gel phase to the liquid phase at 313 K. Calculating the inter-leaflet interaction sides does not reveal significant differences between l-s and l-l systems that could explain the difference in melting temperature. However, we observed, that the lipids in the l-s membranes’ inner leaflet are less ordered compared to the corresponding s-s system. The higher degree of disorder could reflect a higher flexibility of the lipids in the inner leaflet to adapt to the gel structure in the outer leaflet optimizing their interaction and thereby stabilizing the gel phase. However, further research is required to fully understand this difference.

The flip-flop rates measured in the l-l systems were significantly reduced for membranes in the gel phase compared to membranes in the liquid phase at the same concentration of long-chain lipids. From this we conclude, that the gel phase hinders ergosterol from flipping into or out of the gel phase. The diffusion of ergosterol is also reduced, which is in line with the experimental finding of a low sterol content in the gel domains [4]. Additionally, the gel domains could be shown to be stable even when surrounded by lipids in the liquid-ordered phase. Although its size decreases by diffusion of long-chain lipids from the gel phase into the liquid phase, the domain was still intact after several microseconds of simulation time.

This work may set the stage for several natural extensions: (1) To identify the additional impact of the headgroups, comparison with sphingolipids such as IPC, MIPC and M(IP)_2_C may be useful. (2) The flip-flop processes of ergosterol conceptually have different driving forces. First, it reflects the different affinities to different lipids/different phases and, second it may be due to the driving force to adjust the area per lipid in each leaflet in the l-s systems. The additional change of ergosterol population in both leaflets due to the adjustment of the area per lipid as reported in [32] seems to be smaller than the effects seen in this work. Nevertheless, it may be interesting to separate both effects along the lines described in that reference. (3) Although the Martini coarse-grained model has been particularly devised to reproduce lipid behavior [33], a quantitative comparison with experimental results would require the use of atomistic simulations. This holds in particular when comparing parameters like the tilt angle or the melting temperature.

## ACKNOWLEDGMENTS

The authors thank the German Research Foundation for funding (SFB1557-P15 to R. Wedlich-Söldner and SFB1348-A01 to R. Wedlich-Söldner and A. Heuer).

