## Supporting Information for "Effect of very long-chain lipids on the organization of biological membranes: A simulation perspective"

#### Systems

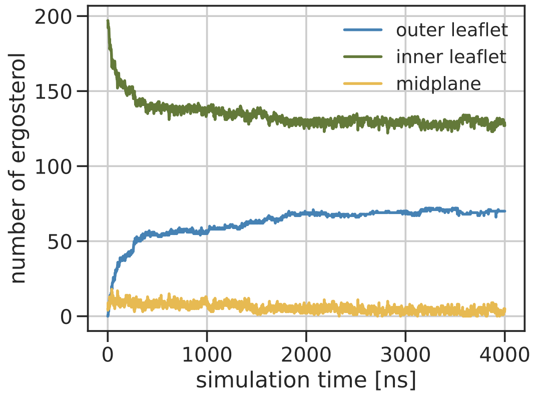


**Figure S1:** The amount of ergosterol in the two leaflets and the midplane during the simulation of the l-s membrane at 313 K with 100 mol-% loPI in the outer leaflet.

#### Analysis methods

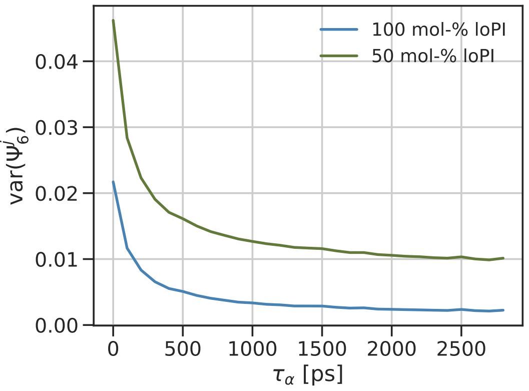


**Figure S2:** For the determination of $\tau_\alpha$, the variance of the hexagonal order parameter is determined for varying values of $\tau_\alpha$ for the l-l membranes with 50 mol-% and 100 mol-% loPI.

### Phase behavior

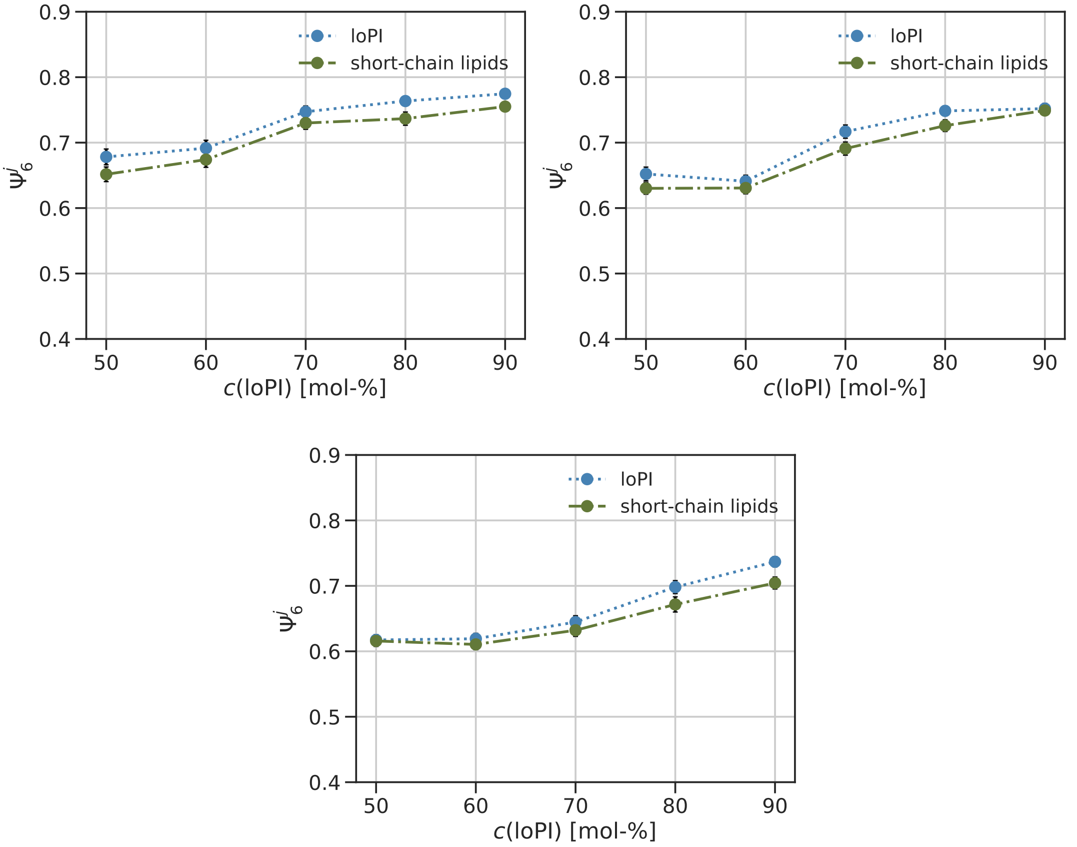


**Figure S3:** The mean hexagonal order parameter for the l-s membranes at 293 K (top left), 303 K (top right), 313 K (bottom) for loPI and the short-chain lipids in the outer leaflet.

### Interdigitation

The interdigitation $I_{\mathrm{eff}}$ is calculated between loPI in the outer and the short-chain phospholipids including ergosterol in the inner leaflet to investigate different contributions to the overall inter-leaflet interactions (figure S4). For the l-s membranes, the trend observed almost matches the trend for all lipids. At lower concentrations, the values are slightly smaller, due to interactions of the short-chain phospholipids in the outer leaflet with the lipids in the inner leaflet (see figure S5). Their contribution decreases as more loPI is added. $I_{\mathrm{eff}}$ is significantly smaller for this lipid selection in the l-l membranes. The contribution of loPI-loPI interactions seems to be dominant here (figure S6). With increasing loPI-concentration, the lipid order of loPI increases leading to an increase in membrane thickness (figure 8 bottom). This causes the other phospholipids to be further apart and fewer interactions are possible. In the l-l membrane in the liquid phase, the interdigitation for loPI-loPI is increasing with the concentration. This might be due to the rising order parameter of the last part of the lipids (see figure 8 right).

While the contributions to $I_{\text{eff}}$ differ between l-l and l-s systems, similar values are obtained. If we assume the interaction strength of loPI with itself and loPI with shorter phospholipids to be similar, the overall inter-leaflet interaction in both systems is the same and can not be the reason for the different melting temperatures. To investigate the cause for this observations, further research is needed.

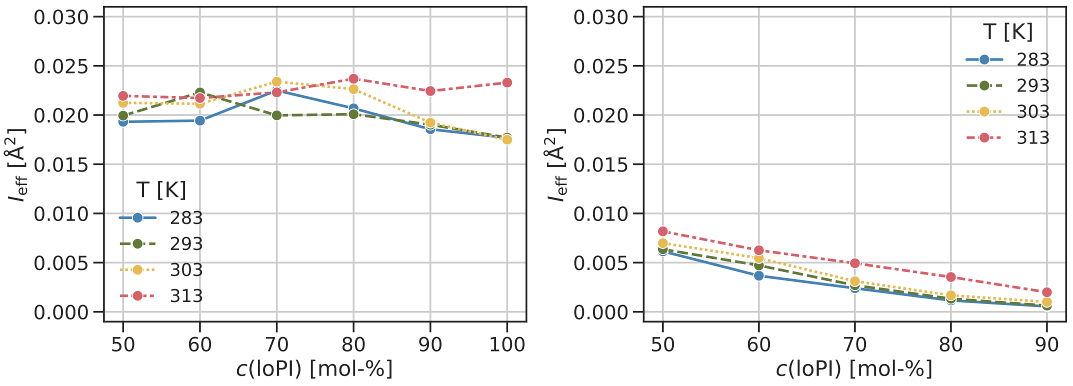


**Figure S4:** The interdigitation $I_{\text{eff}}$ of loPI in the outer leaflet and the phospholipids (not loPI) and ergosterol in the inner leaflet depending on the concentration of loPI at different temperatures in the l-s (left) and l-l membranes (right).

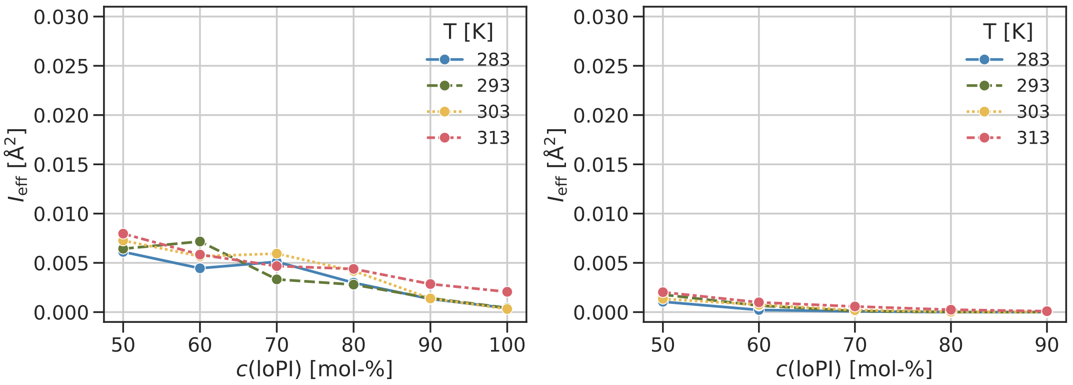


**Figure S5:** The interdigitation $I_{\text{eff}}$ of the phospholipids (excluding loPI) and ergosterol in both leaflets depending on the concentration of loPI at different temperatures in the l-s (left) and l-l membranes (right).

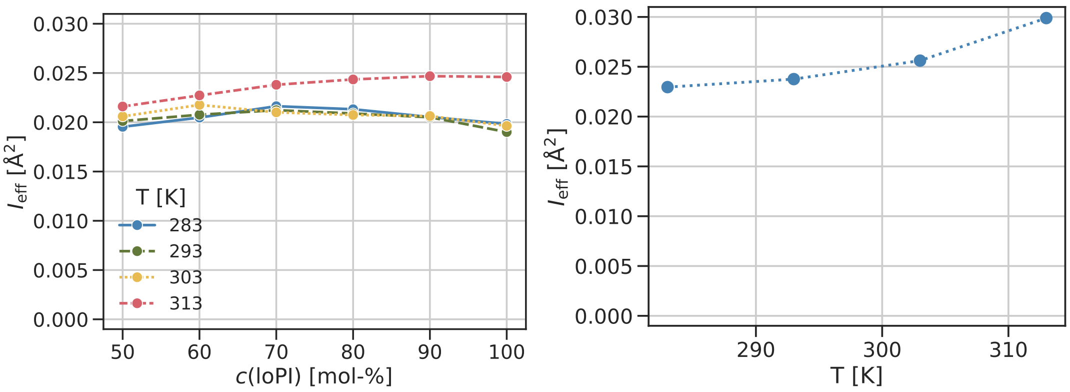


**Figure S6:** The interdigitation $I_{\mathrm{eff}}$ of loPI in both leaflets depending on the concentration of loPI at different temperatures in the l-l membranes (left) and the interdigitation of all phospholipids and ergosterol in the s-s membranes at different temperatures (right).

### Behavior of ergosterol

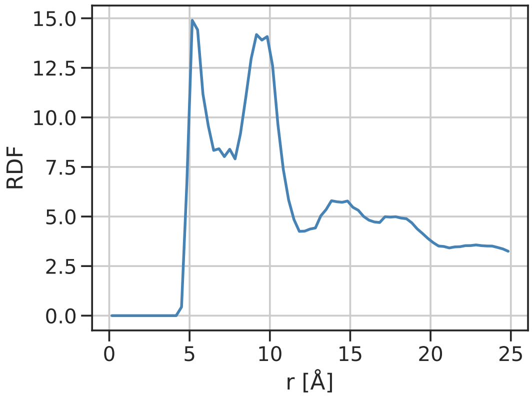


**Figure S7:** The radial distribution function of loPI and POPI PO4 beads in the system with the starting composition of loPI on the left side, ergosterol in the middle and POPI on the right side of the symmetric membrane.

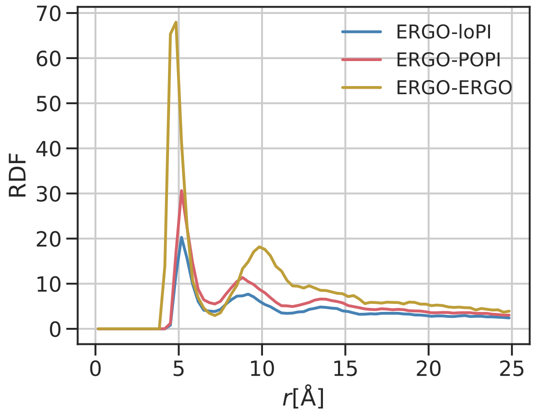


**Figure S8:** Radial distribution function of ergosterol (ROH bead) with loPI, POPI (PO4 beads) and ergosterol in system with the starting composition of loPI on the left side, ergosterol in the middle and POPI on the right side.

### Simulation of a gel domain

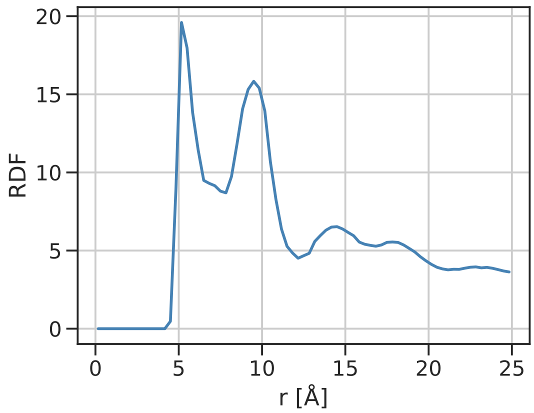


**Figure S9:** The radial distribution function of all phospholipids in the simulation of the gel domain determined at 3500 to 3600 ns simulation time for the PO4 beads.

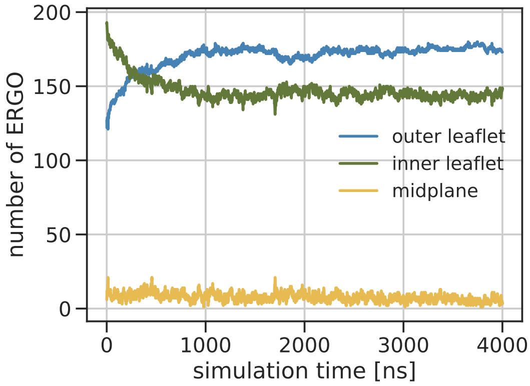


**Figure S10:** Number of ergosterol in the two leaflets and the midplane in the l-s membrane with the gel domain.
